# Extremely-fast construction and querying of compacted and colored de Bruijn graphs with GGCAT

**DOI:** 10.1101/2022.10.24.513174

**Authors:** Andrea Cracco, Alexandru I. Tomescu

## Abstract

Compacted de Bruijn graphs are one of the most fundamental data structures in computational genomics. Colored compacted graphs Bruijn graphs are a variant built on a *collection* of sequences, and associate to each *k*-mer the sequences in which it appears. We present GGCAT, a tool for constructing both types of graphs, based on a new approach merging the *k*-mer counting step with the unitig construction step, and on numerous practical optimizations.

For compacted de Bruijn graph construction, GGCAT achieves speed-ups of 3–21× compared to the state-of-the-art tool Cuttlefish 2 (Khan and Patro, Genome Biology, 2022). When constructing the colored variant, GGCAT achieves speed-ups of 5–39× compared to the state-of-the-art tool BiFrost (Holley and Melsted, Genome Biology, 2020). Additionally, GGCAT is up to 480× faster than BiFrost for batch sequence queries on colored graphs.

## Introduction

De Bruijn graphs are one of the most fundamental data structures in computational genomics, appearing in countless applications, for example, assembly and analysis of sequencing data [10], or RNA-seq data [31], read error correction [19, 33], alignment [21], and compression [2], rearrangement detection [7], just to name a few. To obtain a de Bruijn graph^1^ of order *k* for a multiset of strings (usually sequencing reads, or assembled genomes), for every *k*-mer in the strings, one adds an edge from the node corresponding to its prefix of length *k* − 1, to the node corresponding to its suffix of length *k* − 1. De Bruijn graphs usually have associated also an *abundance* threshold *a*, so that edges (and thus nodes) are added only for *k*-mers appearing at least *a* times in the input strings.

De Bruijn graphs are appealing for several reasons. First, by increasing the abundance threshold, one can have a very simple but effective method of filtering out sequencing errors (i.e., erroneous *k*-mers). Second, having a graph structure allows for a smaller representation of the data in the presence of repeated regions, since equal substrings are represented only once in the graph. Third, by focusing on maximal *non-branching* paths, i.e., maximal paths whose internal nodes have indegree and out-degree equal to 1 (also called *maximal unitigs*), one can discover “variation-free” regions.

Originally, maximal unitigs were introduced for the genome assembly problem; for example, assembly contigs are usually unitigs of a corrected assembly graph [12, 32]. However, by replacing each unitig with e.g., an edge labeled with the label of the unitig (label obtained by identifying the overlapping (*k* −2)-mers), one gets an equivalent graph, but of much smaller size. Such a graph is usually called a *compacted de Bruijn graph*.^2^ Many of the applications cited above actually use a compacted graph, due to their equivalent representation power, but significantly smaller size. In fact, if one just wants to represent the *k*-mers of a dataset in plain text form, there exist more efficient equivalent representations [29, 6, 16] (and even optimal [35, 34]), but all of these use maximal unitigs as a starting point.

A popular variant of a de Bruijn graph is the *colored* de Bruijn graph, originally introduced for de novo assembly and genotyping of variants [15]. Such graph is built from a *collection* of datasets, for example different sequencing datasets or different (full) genome sequences. For every *k*-mer, colored de Bruijn graphs also store the identifiers (*colors*) of the datasets in which the *k*-mer appears. One can think of a colored de Bruijn graph as a compressed representation of the *k*-mers in a *collection* of datasets, but which retains enough information (i.e., the color of every *k*-mer) in order to identify each dataset in this combined graph. Later applications include pangenomics [37], RNA-seq quantification [5], bacterial genome querying [22], alignment and reference-free phylogenomics [37], microbiome research [13], just to name a few.

While computing the maximal unitigs of a de Bruin graph can be done with a simple linear-time algorithm, the practical hardness of the problem stems from the fact that the initial de Bruijn graph does not fit the main memory in applications. Practical tools computing compacted de Bruijn graphs have to cleverly use disk to store intermediary data, partition the data in order to use many CPU cores efficiently, and minimize CPU, RAM and I/O bottlenecks. One of the first major tools for computing maximal unitigs was BCALM2 [9]. BCALM2 first does a *k*-mer counting step inspired by KMC2 [11] and a filtering pass based on the multiplicity of the *k*-mers. Then it finds *k*-mers that should be joined together by bucketing them on their left/right minimizers [30] (corresponding to the minimizers of the leftmost and rightmost (*k* − 1)-mers). Each bucket is processed independently and in parallel to find all the possible extensions. Finally, BCALM2 glues all the unitigs that were produced in different buckets together using a union-find data structure. The state-of-the-art tool for maximal unitig computation is Cuttlefish 2 [16]. It starts with an initial *k*-mer counting step using KMC3 [18]. It employs a perfect hash computation on the *k*-mers using BBHash [20]. The key insight relies on a novel automaton-based approach to compute the branching state of every (*k* − 1)-mer, using only the minimum amount of information, i.e., zero, one, or more than one left / right neighbors. Then, it builds the graph by looking at the automaton of every (*k* − 1)-mer, extending the unitig if the current (*k* − 1)-mer does not branch forwards and the following (*k* − 1)-mer does not branch backwards. Cuttlefish 2 tends to significantly write to disk to further resplit the intermediate buckets and keep the maximum memory usage low. Like BCALM2, it also does not compress the buckets on disk (except for prefix collapsing) and thus very repetitive datasets still require large disk I/O. Moreover, for higher *k* values, KMC3 tends to use more time and memory, as it has to store the exact *k*-mers all the time, which in consequence also affects the behavior of Cuttlefish 2.

The state-of-the-art tool for maximal unitig computation with associated color information is BiFrost [14]. It uses an in-memory only approach, with various blocked Bloom filters [4, 28] partially indexed by minimizers that approximate the *k*-mers present in the final graph, and does several passes on the original input to remove the false edges wrongly created due to the use of Bloom filters. Then it internally stores the *k*-mers grouped by minimizers, allowing for relatively fast deletions and insertions of new *k*-mers. However, while blocked Bloom filters are very memory efficient, they are also not very cache efficient (even with the infra-block sse2 optimizations done in BiFrost). Also the memory representation of the *k*-mers gives a tradeoff between the ease of doing small updates to the graph and the speed of inserting batches of *k*-mers, thus the build time of the graph is still considerably high. To (optionally) build a colored graph, it uses various types of compressed bitmaps (roaring [8], or simple bitsets) to store the set of colors of each *k*-mer. While this allows fast insertions and querying, it stores redundant color sets information since *k*-mers that share the same set of colors are still encoded as two separate sets.

## Results

### GGCAT overview

We propose a new tool for constructing compacted, and optionally colored, de Bruijn graphs, GGCAT. As opposed to BCALM2 and Cuttlefish 2, the first idea of GGCAT is to merge the *k*-mer counting step with unitig construction, by adding a little more “context” information that allows us to compute valid global unitigs inside each bucket that the input is split into. This avoids the storage of every single *k*-mer, since only unitigs built inside the buckets are written to disk. Moreover, as opposed to other tools, these unitigs are lz4-compressed before writing to disk, which allows for a substantial reduction in disk usage for highly repetitive datasets. Second, we avoid a union-find data structure (used by BCALM2) with a new joining step across buckets, that guarantees exact results with very low *expected* running time. Third, we devise a parallelization pipeline that divides the algorithm into smaller execution units (e.g., reading from disk, *k*-mer counting, *k*-mer extension), thus preventing core stalling due to waiting for data.

On the theoretical side, we give a string-based definition of maximal unitig in the presence of reverse complements (*canonical maximal unitig*, Definition 1) that (i) allows us to avoid introducing a heavy formalism based on e.g. bidirected de Bruijn graphs and (ii) closely mimics our algorithm, thus leading to a simple proof of correctness. Moreover, since our unitigs are in an edge-centric graph, in the Supplemental Material we prove that they are equivalent to node-centric unitigs in a node-centric graph, as employed by e.g. BCALM2 [9] (and which we also confirm experimentally), result which we did not find in the literature and may be of independent interest.

For colored graphs we extend our algorithm above with an approach inspired by BiFrost, but with several optimizations that allow comparable colormap sizes with substantially improved build times. The main difference w.r.t. BiFrost is that, instead of using an individual (compressed) color bitmap for each possible *k*-mer, GGCAT maps each color set to a *color set index*, approach similar to e.g. [1, 27, 23]. Moreover, to store each color set, we compute the difference between consecutive colors and compress them using a run-length encoding. Finally, when storing to disk, the color set indices of the consecutive *k*-mers of each unitig are also run-length encoded. This strategy proves efficient because unitigs are “variation-free”, and thus usually have few color set indices associated to their *k*-mers. Since CuttleFish 2 is significantly faster than BiFrost (on non-colored graphs), these ideas, combined with our improvements over Cuttlefish 2, leads to a major speed up over BiFrost for colored graphs.

Similarly to BiFrost [14], GGCAT also supports querying the produced colored graph against batch input sequence queries. More precisely, for every query sequence, in the uncolored case we need to return the number (equivalently, percentage) of *k*-mers of the sequence that also appear in the entire target graph. In the colored case, for every color *c*, we need to return the number of *k*-mers of the query sequence matching *k*-mers of the graph that are colored with *c*. In practice, we need to query many input sequences at the same time (e.g., a fasta file). GGCAT solves both types of batch queries by an approach very similar to the graph construction steps.

Finally, being written in Rust, GGCAT also integrates the matchtigs [34] and eulertigs [35] Rust libraries, (optionally) computing thus minimum plain-text representations of the set of *k*-mers of the input fasta files.

### Tested tools, datasets and hardware

To compute compacted de Bruijn graphs, we chose to compare only against Cuttlefish 2, since the article introducing it [16] showed that it significantly outperforms popular tools such as BCALM2 [9] and BiFrost [14] (in its non-colored variant). To compute colored de Bruijn graphs, we chose to compare only against BiFrost, since the article introducing it [14] showed that it significantly outperforms popular tools such as VARI-merge [26]. We decided to not compare against Cuttlefish 1 [17] for colored graphs because it adopts a different convention for colors (each unitig can have only one subset of colors) and does not support querying the resulting graph. We run all tools in their default settings (see the Supplemental Material for the commands used).

For *k* ≤ 64 GGCAT represents *k*-mers exactly. Also, to support values larger than 64, GGCAT uses a non-bijective 128-bit Rabin-Karp hash function to represent each *k*-mer (where each of the four bases is represented by a different prime number), to avoid storing it in full length. In extremely rare cases, it can lead to some collisions in hash values, that can cause unwanted joining of some unitigs or extra splittings of a maximal unitig. GGCAT can detect (but not correct) most of the collisions, warning the user if some errors can be expected in the graph. In all the tested datasets with *k >* 64, we found no occurrence of a hash collision.

For the uncolored case, we use an Illumina whole-genome sequencing Human read dataset, a Human gut microbiome read dataset, 309K Salmonella genome sequences, and 649K Bacterial genomes. For the colored case, we use 100 Human genome sequences, 100K Salmonella sequences from the full 309K Salmonella dataset (to save computational resources), and all Bacterial genomes. See Sec. Data access for accession details, and Table 1 with structural characteristics of these datasets. For the read datasets we use an abundance threshold of 2, and for the genome reference datasets we use an abundance threshold of 1. For a sanity check, we checked that for GGCAT the maximal canonical unitigs are exactly equivalent to the ones produced by BCALM2, for the uncolored graphs produced from 1K Salmonella genomes, and from the Human read dataset.

**Table 1.**
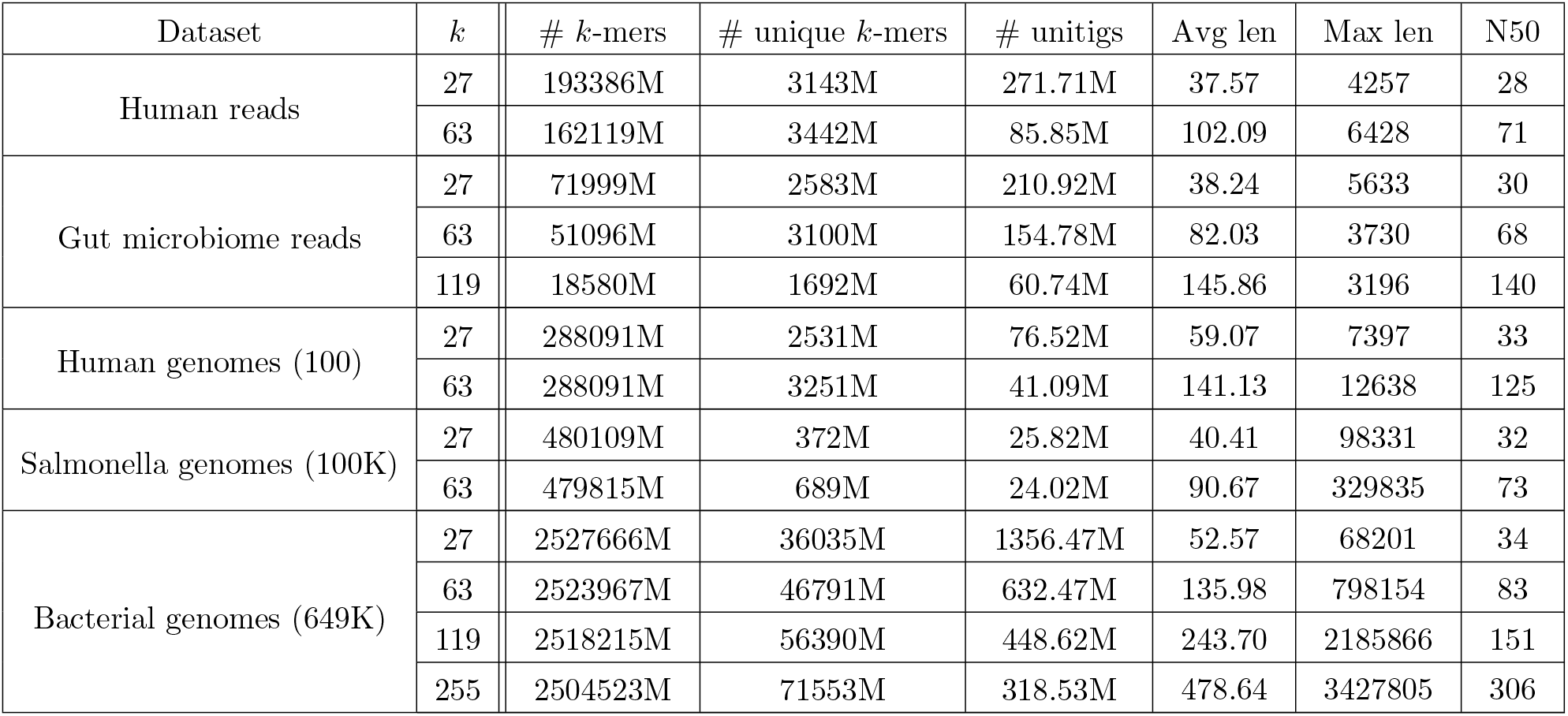
Datasets statistics with various values of *k*. The first column considers the whole input dataset, while the rest of the columns refer to the graph built from the dataset with the specified *k* value. The Avg, Max and N50 columns are computed on the lengths of the maximal unitigs of the graph.

We ran the experiments on three servers of increasing power: a *small* server with an AMD Ryzen 5 3600 6-core CPU, 64GB RAM and a RAID 0 with two 7200RPM HDDs; a *medium* server with an AMD Ryzen Threadripper PRO 3975WX 32-core CPU, 512GB RAM and a RAID 5 7200RPM HDD; a *large* server with two AMD EPYC 7H12 64-Core CPUs, 2TB RAM and a SATA SSD.

### Construction results

In the uncolored case, for the Human read dataset, we run two realistic values of *k*, 27 and 63. On the other three datatsets we tested the behavior for larger *k* values, where the graphs still remain complex: for Gut microbiome reads, *k* = 119, and for the 309K Salmonella genomes and 649K Bacterial genomes, *k* ∈ {119, 255}. To save computational resources, we did not run Cuttlefish 2 for the latter dataset for larger *k* values, since the tool does not scale. We show the results in Table 2.

**Table 2.** Uncolored construction, wall clock time and memory usage. The Bacterial test was performed on the large server with 16 threads while the other tests on the small server with 12 threads.

| Dataset | $k$ | Cuttlefish 2 | GGCAT |
| --- | --- | --- | --- |
| Human reads | 27 | 1h:15m (3.95GB) | 1h:16m (4.54GB) |
|  | 63 | 2h:07m (4.23GB) | 1h:03m (7.11GB) |
| Gut microbiome reads | 27 | 0h:30m (3.35GB) | 0h:22m (6.09GB) |
|  | 63 | 1h:08m (3.86GB) | 0h:19m (5.42GB) |
|  | 119 | 1h:04m (3.13GB) | 0h:12m (5.33GB) |
| Salmonella genomes (309K) | 27 | 6h:59m (4.38GB) | 3h:38m (3.46GB) |
|  | 63 | 12h:02m (3.88GB) | 3h:31m (3.96GB) |
|  | 119 | 17h:07m (3.95GB) | 3h:39m (4.12GB) |
|  | 255 | 77h:58m (4.82GB) | 3h:44m (4.33GB) |
| Bacterial genomes (649K) | 27 | 13h:24m (42.60GB) | 7h:7m (8.49GB) |
|  | 63 | 25h:29m (53.84GB) | 5h:56m (10.06GB) |
|  | 119 | – | 6h:14m (9.46GB) |
|  | 255 | – | 6h:30m (9.42GB) |

For Human reads and *k* = 27, GGCAT has a similar performance as Cuttlefish 2. However, forlarger *k* values, and on the other three larger datasets, GGCAT outperforms Cuttlefish 2 in terms of speed by up to 4.3×, for *k* ≤ 63. The main speed improvements come from our read splitting strategy, which proves most useful for larger *k* values. For larger *k* values (119 and 255), GGCAT is faster than Cuttlefish 2 by up to 20.8×. (Notice also that, as opposed to GGCAT, Cuttlefish 2 does not support *k* values larger than 255.) Despite this, in all cases GGCAT has an overall memory usage in the same order of magnitude as Cuttlefish 2, or even substantially lower in the complex Bacterial genomes dataset.

We also tested the scalability of GGCAT by computing the uncolored graph of an increasing number of Salmonella genomes (see the Supplemental Material), the results showing a linear relation between the number of genomes and the running time.

The colored construction results are in Table 3. Compared to BiFrost, in the first dataset GGCAT is 5.1 times faster for *k* = 27 and 4.6 times faster for *k* = 63. For the Salmonella genomes, for *k* = 27, GGCAT is 33.3 times faster than BiFrost, and for *k* = 63 GGCAT is 39.3 times faster than BiFrost. For *k* = 27 BiFrost craqshed, while for *k* = 63 we stopped its run after 10 days to save computational resources. Instead, GGCAT completed in both cases in under 14 hours. The memory used by GGCAT in the colored construction tests is from 3.7 to 12 times less than BiFrost, but this is not directly comparable since GGCAT uses disk intermediate storage, while BiFrost uses a fully in-memory algorithm.

**Table 3.** Colored construction, wall clock time and memory usage, using 16 threads. The Human and Bacterial benchmarks were executed on the large server and the Salmonella benchmark on the medium server.

| Dataset | $k$ | BiFrost colored | GGCAT colored |
| --- | --- | --- | --- |
| Human genomes (100) | 27 | 6h:39m (30.68GB) | 1h:17m (8.29GB) |
|  | 63 | 5h:16m (38.65GB) | 1h:08m (8.80GB) |
| Salmonella genomes (100K) | 27 | 50h:29m (79.32GB) | 1h:31m (6.54GB) |
|  | 63 | 46h:34m (85.21GB) | 1h:11m (6.18GB) |
| Bacterial genomes (649K) | 27 | *crashed* | 13h:28m (21.64GB) |
|  | 63 | *> 10 days* | 11h:33m (21.48GB) |

### Colored querying results

To test querying, we used the colored graphs of 100 Human genomes for *k* ∈ {27, 63} produced in the previous test, and queried them using 4 million 250bp sequencing reads from the first dataset of Human reads. Results (in Table 4) show that GGCAT outperformed BiFrost by 83.6 times for *k* = 63, while for *k* = 27 GGCAT was more than 480 times faster than BiFrost. This significant improvement is due to the fact that we implement querying as a natural extension of the unitig construction step, benefiting thus from all its optimizations.

**Table 4.** Querying in the colored graph of 100 Human genomes, wall clock time and memory usage, using 16 threads.

| Query | $k$ | BiFrost | GGCAT |
| --- | --- | --- | --- |
| 4 million reads | 27 | 33h:14m (30.72GB) | 4m:05s (17.48GB) |
|  | 63 | 5h:26m (35.21GB) | 3m:54s (11.10GB) |

## Discussion

Computing a compacted de Bruijn graph (and optionally colored) is one of the most fundamental problems in computational genomics, with a long history of computational tools developed for this problem. GGCAT pushes the boundary in terms of a highly efficient implementation, based on both novel algorithmic aspects (e.g. combining *k*-mer counting with unitig construction, and a new strategy of joining partial unitig across buckets), but also in terms of an efficient parallelization pipeline minimizing idle CPU cores and disk I/O bottlenecks, or further optimizations such as lz4-compression of data written to disk. GGCAT reduces sequence queries to an approach similar to graph construction, benefiting thus from its highly optimized architecture.

Overall, this leads to a several times improvement over Cuttlefish 2 (and even bigger for larger *k* values), and a two orders of magnitude improvement over BiFrost for colored construction and querying. GGCAT can thus have a significant impact in all downstream analyses that require computing a compacted de Bruijn graph.

## Methods

### Preliminaries

In this paper all strings are over the same alphabet ∑ = *{A, C, G, T }*. We denote concatenation of two strings *x* and *y* as *x · y*. If *x* is a substring of *y*, we write *x* ∈ *y*. We denote the length of a string *x* as |*x*|. Given a string *x* of length at least *k*, we denote by pre_*k*_ (*x*) the prefix of *x* of length *k*, and by suf_*k*_(*x*) the suffix of *x* of length *k*. For two strings *x* and *y* such that suf_*k*_(*x*) = pre_*k*_ (*y*), we denote by *x*⊙ _*k*_ *y* the string *x ·* suf_|*y*|−*k*_ (*y*) (the *merge* of *x* and *y*). Given a set *S*, we denote ends_*k*_(*S*) = *{*pre_*k*_ (*x*), suf_*k*_(*x*) : *x* ∈ *S}*.

A *k-mer* is a string of a given length *k* over the alphabet ∑. Given a *k*-mer *q*, we say that *q* is a *k*-mer of a string *x* if *q* ∈ *x* (in this case, we also say that *q appears*, or *occurs* in *x*). Given a set or a multiset *S* of strings, we say that *q appears* in *S*, and write *q* ∈ *S*, if *q* appears in some string in *S*. The *edge-centric* de Bruijn graph or order *k* of a multiset *R* of strings is defined as the directed graph having as nodes the (*k* − 1)-mers appearing in *R*, where we add an edge from a node *x* to a node *y* if suf_*k*−2_(*x*) = pre_*k*−2_(*y*) and *x* ⊙_*k*−2_ *y* ∈ *R*. That is, the set of edges is exactly the set of all *k*-mers of *R*. In such an edge-centric graph, the *spelling* of a path *P* = (*x*_1_, …, *x*_*t*_) is the string *x*_1_ ⊙_*k*−2_ *x*_2_ ⊙_*k*−2_ *· · ·* _*k*−2_ *x*_*t*_.

A *unitig*^3^ is defined as a path^4^ of the de Bruijn graph of *R* containing at least one edge, such that all the internal nodes of the path (i.e., different from the first and the last) have in-degree and out-degree equal to one. A *maximal* unitig is one that cannot be extended by one node without losing the property that it is a unitig. We are interested in outputting the *set* of all maximal unitigs of a de Bruijn graph, where for unitigs that are cycles we need to output only one cyclic shift (i.e., in the output, there can be no equivalent cycles). For ease of notation, by unitig we will also refer to its *spelling*. Clearly, the first and last node of a unitig different from a cycle must satisfy the condition that *either* its in-degree is different from one, or its out-degree is different from one. Note that under this definition, the maximal unitigs form a partition of the edges, i.e., of the *k*-mers of *R*.

For the rest of this paper, we consider an alternative definition of maximal unitigs that does not explicitly use a de Bruijn graph. This has several advantages: it connects to the recent literature on *spectrum preserving string sets* (unitigs being one such type of sets) [29, 6, 34], it naturally extends to reverse complements without introducing heavy definitions related to bidirected de Bruijn graphs, and ultimately it matches our algorithm, which proceeds bottom-up by iteratively merging *k*-mers and unitigs as long as possible (i.e., the existence of branches in the de Bruijn graph is checked implicitly via *k*-mer queries).

Given a multiset *S* of strings, and a string *x*, we denote by occ(*x, S*) the number of occurrences of *x* in the strings of *S*, each different occurrence in a same string in *S* being counted individually. Given a string *x* ∈ ∑, we denote by *x*^−1^ ∈ ∑ the reverse complement of *x*. Given a multiset *S* of strings, and a string *x*, if *x* ≠ *x*^−1^, we define occ_rc_(*x, S*) = occ(*x, S*) + occ(*x*^−1^, *S*), otherwise occ_rc_(*x, S*) = occ(*x, S*). We analogously define app(*x, S*) = min(1, occ(*x, S*)) and app_rc_(*x, S*) = min(1, occ_rc_(*x, S*)).

Given multisets of strings *R* and *U*, we say that *R* and *U* have the same *k-mer spectrum* if any *k*-mer that appears in one of the sets also appears in the other set. Analogously, we say that *R* and *U* have the same *canonical k-mer spectrum* if any *k*-mer *q* that appears in one of the sets, *q* or *q*^−1^ appears in the other set. We can equivalently express the fact that sets *R* and *U* have the same non-canonical *k*-mer spectrum with the condition:

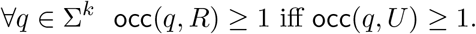

Likewise, *R* and *U* have the same *canonical k-mer spectrum* if:

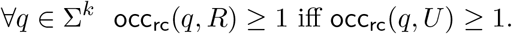

We now give an equivalent *string-centric* definition of the *set U* of maximal unitigs of a multiset *R* of strings, under our formalism. As a warm-up, we start with the case when we do not have reverse complements.

First, we require that all strings in the *set U* have length at least *k*, meaning unitigs contain at least one edge. Second, we require that *R* and *U* have the same *k*-mer spectrum. Third, if a (*k* − 1)-mer appears at least two times in *U*, then it cannot be an internal node in any unitig.

In other words, we forbid merging two separate unitigs at a branching (*k* − 1)-mer, since such branching (*k* − 1)-mer must appear in at least one other string in *U* :

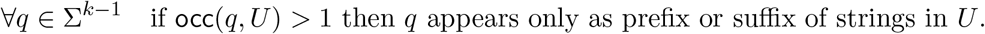

Note that the above property also ensure that no two equivalent cyclic unitigs are in *U*.

To impose also maximality, we state that a (*k* − 1)-mer is a prefix or a suffix of a unitig if and only if it is either branching, a sink, or a source:

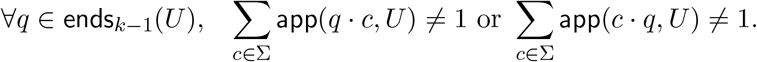

Having defined maximal unitigs without reverse complements, we now give the string-centric definition of maximal unitigs assuming also reverse complements (which we call *canonical maximal unitigs*). In fact, we give a more general one, handling also a required abundance threshold of the *k*-mers in *R*, further underlining the flexibility of our string-centric view.

#### Definition 1

(Canonical maximal unitigs). *Given a multiset R of strings and integers k* ≥ 2 *and a* ≥ 1, *we say that a* set *U of strings is the set of* canonical maximal unitigs *of R with k-mer size k and abundance threshold a if the following conditions hold:*

1. ∀*x* ∈ *U*, |*x*| ≥ *k, and* ∀*x, y* ∈ *U, x* ≠ *y*^−1^ *(note that x* ≠ *y is guaranteed by the fact that U is a set);*
2. ∀*q* ∈ ∑^*k*^ occ_rc_(*q, R*) ≥ *a iff* occ_rc_(*q, U*) ≥ 1 *(same canonical k-mer spectrum, with abun-dances);*
3. ∀*q* ∈ ∑^*k*−1^, *if* occ_rc_(*q, U*) *>* 1 *then q and q*^−1^ *appear only as prefix or suffix of strings in U (unitigs do not span over branching* (*k* − 1)*-mers);*
4. ∀*q* ∈ ends_*k*−1_(*U*), _*c*∈∑_ app_rc_(*q · c, U*) ≠ 1 *or* _*c*∈∑_ app_rc_(*c · q, U*) ≠ 1 *(maximality)*.

In our algorithm, we build unitigs incrementally, starting from individual *k*-mers (i.e., individual edges of the de Bruijn graph, which are unitigs), and extending them in both directions, as long as the resulting string remains a unitig (by checking for the satisfaction of condition 3 in Definition 1 at each step, i.e. whether we have reached the end of a unitig or not). Even though this is a simple strategy, behind other tools such as [9], it is non-trivial how to implement this efficiently in terms of running time, memory consumption, disk usage, and parallelization.

Given an integer *m* ≤ *k*, and a rolling hash function hash : ∑^*m*^ → ℤ, the *minimizer* of a *k*-mer *x* is

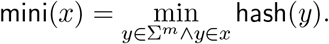

Note that in this definition the minimizer is only a hash value, and does not keep track of the particular position of the *m*-mer that has that minimum hash value.

Throughout the algorithm, we will refer to *buckets* or *groups* as a partition of the data that is stored as a single blob, for example when stored on disk each bucket corresponds to a file. Multiple buckets are used to partition data in a way that is optimized for parallelization, thereby allowing for parallel and independent processing of each bucket. They are also used to reduce the memory consumption of the algorithm, since only the buckets that are currently being processed occupy main memory, while the other ones use only disk space.

In the rest of this section we present the algorithm for maximal unitigs without reverse complements, and then in Sec. Construction correctness we explain the changes for canonical maximal unitigs.

### Read splitting

Each read *R*_*j*_ is split into substrings 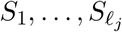 that overlap on *k* − 2 characters, such that all (*k* − 1)-mers of *S*_*i*_ have the same minimizer, for all *i* ∈ *{*1, …, *ℓ*_*j*_*}*. For the minimizer hash function hash we use the ntHash [25] function, because it can give fast computation while ensuring good randomness in its value. Note that we can have multiple minimizer locations in the same substring *S*_*i*_ as long as they have the same hash value. We can compute 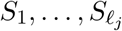 in linear time in the size of *R*_*j*_ as follows. First, for every *m*-mer *x* of *R*_*j*_, we compute hash(*x*) in a rolling manner. Then, in a sliding window manner, we compute the minimum of each window of *k* − *m* consecutive *m*-mers (which correspond to a (*k* − 1)-mer). Finally, we group consecutive (*k* − 1)-mers that share the same minimum in their corresponding window. For efficiency, we perform these three steps in a single pass over *R*_*j*_.

For every *S*_*i*_ obtained in this manner, let *a* and *b* be the characters of *R*_*j*_ immediately preceding and succeeding *S*_*i*_ in *R*_*j*_, respectively, or $ if they do not exist. We call *a* and *b linking characters*. Consider the string 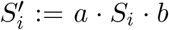 and observe that 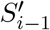 and 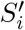 have a suffix-prefix overlap of *k* characters, since *S*_*i*−1_ and *S*_*i*_ have a suffix-prefix overlap of *k* − 2 characters and we added *b* at the end of *S*_*i*−1_ and *a* at the beginning of *S*_*i*_. Recall that all (*k* − 1)-mers of *S*_*i*_ have the same minimizer, say *h*; we assign each extended string 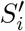 to a *group G*_*h*_ associated to such unique minimizer *h*. We say that a *k*-mer *x* appears in a group *G*_*h*_ if *x* is a substring of some 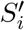 in *G*_*h*_. See Figure 1 for an illustration.

**Figure 1:**
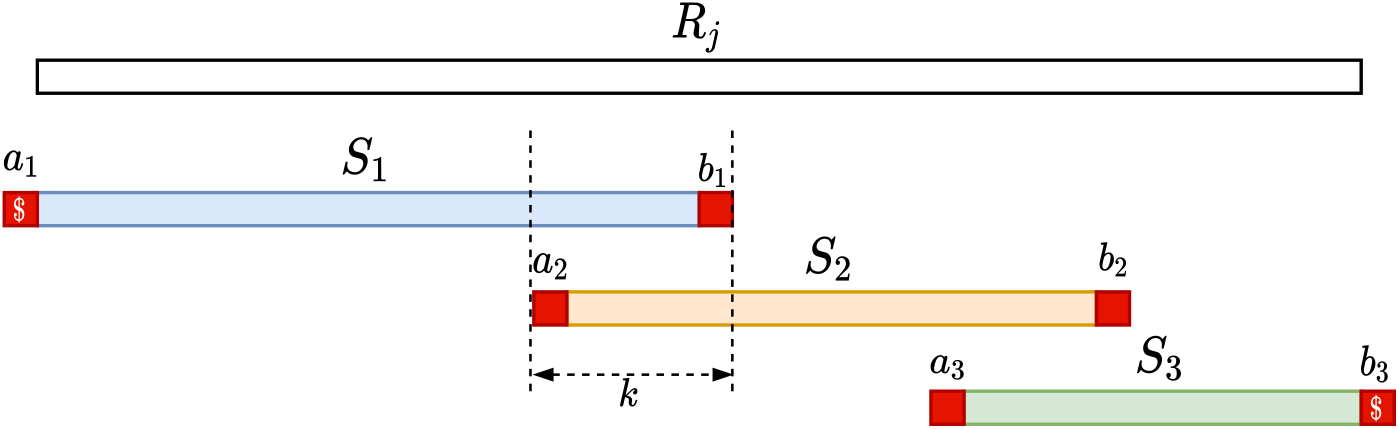
Illustration of the read splitting step. Read *R*_*j*_ is split into substrings *S*_1_, *S*_2_, *S*_3_ such that all *k*-mers of each *S*_*i*_ have the same minimizer, and extra linking characters (in red) are added to each *S*_*i*_. The overlap between two such consecutive extended *S*_*i*_’s is of exactly *k* characters.

The above grouping strategy is similar to the one in [18], applied to *k*-mers instead of (*k* − 1)-mers (our strings 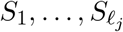 are called *super* (*k* − 1)*-mers* in [11]), with the exception that when we group we add the linking characters. The following simple properties are key to ensuring the correctness of our approach.

#### Lemma 1.

*Let x be a k-mer appearing in the reads, and in a group G*_*h*_. *The following properties hold:*

a. *There is at most one other group in which x appears, and moreover, x appears in two distinct groups if and only if* mini(pre_*k*−1_(*x*)) ≠ mini(suf_*k*−1_(*x*)).
b. occ(*x, G*_*h*_) = occ(*x, R*);
c. *If* mini(*x*) = *h (i*.*e*., *x does not contain a linking character), and it has a* (*k* − 1)*-suffix-prefix overlap with some k-mer y (in either order), then also y appears in group G*_*h*_.

*Proof*.

a. (a) Any *k*-mer has only two (*k* − 1)-mers, which in the worst case have different minimizers *h*_1_ and *h*_2_, and as such *x* can appear in at most two groups 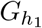 and 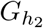, and these are different if *h*_1_ ≠ *h*_2_.
b. This follows by construction of the group, since all the *k*-mer occurrences that have the same minimizer are put in the same group.
c. If also the minimizer of *y* is *h* (i.e., it does not contain a linking character), then also *y* appears in *G*_*h*_. If not, recall that we added an extra character at the beginning and end of every string assigned to *G*_*h*_, thus *y* is a *k*-mer containing a linking character and thus appears in *G*_*h*_.

Since we want to write the groups to disk, and their number is the number of distinct minimizers, we merge the groups into a smaller number of buckets, that are written to disk.

### Construction of intermediate unitigs

Lemma 1 ensures that extending any *k*-mer *x* can be correctly performed just by querying the group of *x*.

For each group, we perform:

1. A *k*-mer counting step of the strings in the group, using a hashmap, while also keeping track if a *k*-mer contains a linking character. More precisely, we scan each string in a group, and for each *k*-mer that we encounter we increase by one its abundance in the hashmap, and add a flag it if contains a linking character.
2. From the hashmap, we create a list of unique *k*-mers of the group, that have the required abundance. This abundance check is correct thanks to Lemma 1(b).
3. We traverse the list of *k*-mers, and for each non-used *k*-mer *x*, we initialize a string *z* := *x*, which will be extended right and left as long as it is a unitig (see Figures 2 and 3). We try to extend *z* to the right by querying the hashmap for suf_*k*−1_(*z*) *· c*, for all *c* ∈ *{A, C, G, T }*. If there is a unique extension *y* such that suf_*k*−1_(*z*) = pre_*k*−1_(*y*), then we query the hashmap for *c ·* pre_*k*−1_(*y*), for all *c* ∈ *{A, C, G, T }*. If exactly one match is found (i.e., suf_*k*_(*z*)), then we replace *z* with *z* _*k*−1_ *y*, and we mark *k*-mer *y* as used in the hashmap. If *y* is not marked in the hashmap as having a linking character, then we repeat this right extension with the new string *z*. The queries to the hashmap are correct thanks to Lemma 1(c). When we stop the right extension, we perform a symmetric left-extension of *z*. After both extensions are completed, the resulting unitig *z* is given a unique index *id*_*z*_. If the extension of *z* was stopped because of a linking character in the first or last *k*-mer *y* of *z*, we add (*y, id*_*z*_) to a list *L*.

**Figure 2:**
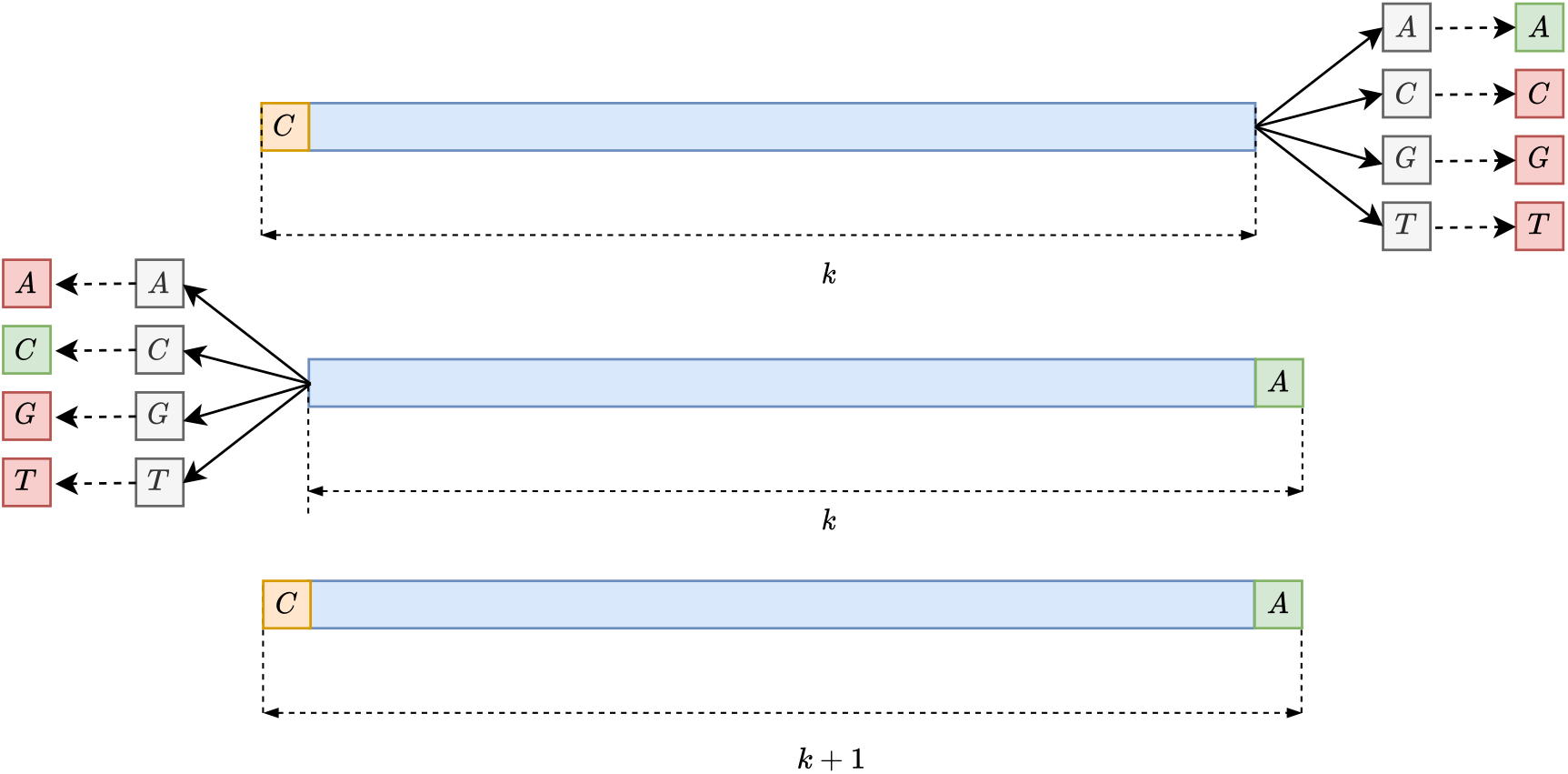
The extension step of the intermediate unitig construction happens inside each group. For each *k*-mer (top) it looks for a possible extension by checking all the 4 possible neighbor *k*-mers in both direction, and extends the *k*-mer (bottom) only if there is exactly one match both forwards and backwards (depicted in green in the first two figures from the top). Then it repeats the same process until no more extensions can be performed.

**Figure 3:**
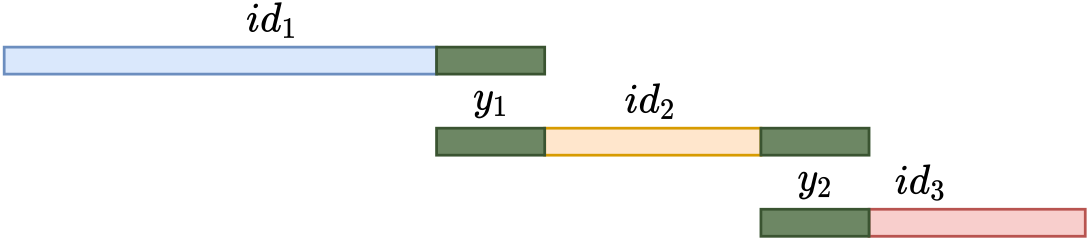
The result of the intermediate unitig construction. Each intermediate unitig that has a possible extension shares an ending with another intermediate unitig.

Notice that, after all groups have been processed, for any (*y, id*_*z*_) in *L*, there exist exactly one other (*y, id*_*z*_*′*) in *L*, added from a different group, by Lemma 1(a). These two tuples indicate unitigs that have to be iteratively merged to obtain the maximal unitigs.

### Unitig merging

The tuples (*y, id*_*z*_) in *L* are sorted by *y*, such that the two entries (*y, id*_*z*_) and (*y, id*_*z*_′) appear consecutively. Moreover, for any unitig *z*, there are at most two entries (*x, id*_*z*_) and (*y, id*_*z*_) in *L* (corresponding to its two endpoints). From these, we construct a list (*id*_*z*_, *id*_*z*_′) that is put in another list *P* of pairs of unitig that must be merged into maximal unitigs. This is one of the hardest steps to parallelize, since no partitioning can be done ahead of time that puts all the unitigs that are contained in a maximal unitig in the same partition. In other tools, e.g., BCALM2 [9], this step is done using an union-find data structure, that can be difficult to be used with concurrency. Our solution uses a randomized approach (i.e., with guaranteed correctness and only *expected* running time) to put in the same partition the unitigs that should be merged, repeating the process until all the unitigs are merged into the final maximal unitigs (see Figure 4).

**Figure 4:**
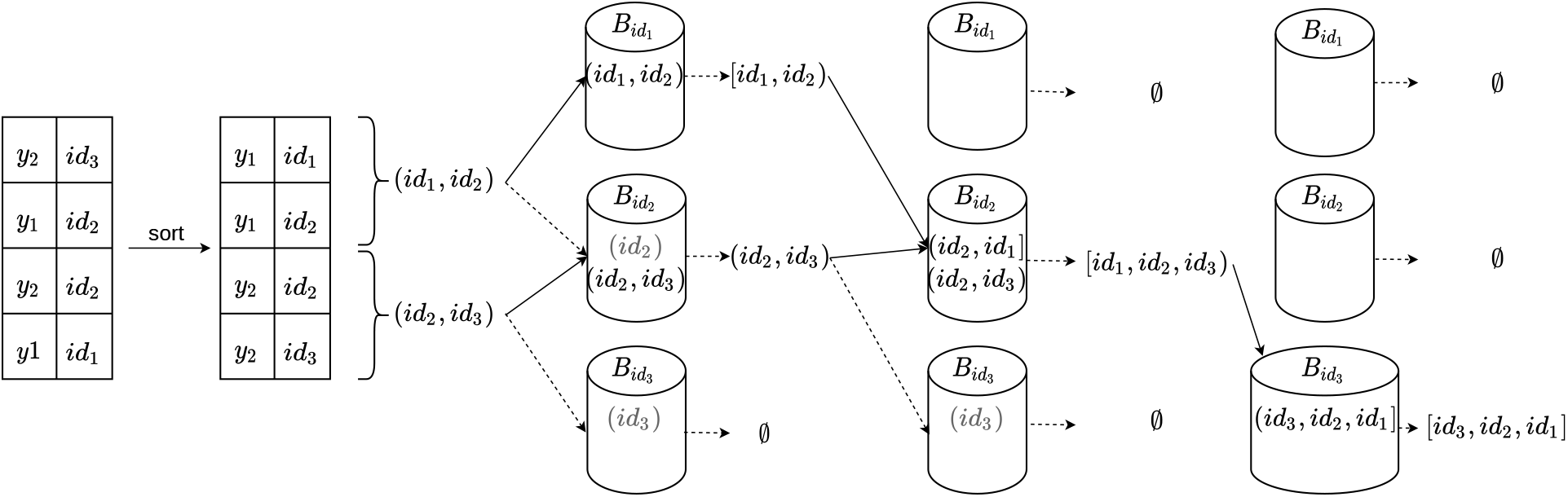
The unitig merging step on the unitigs from Figure 3. Each pair (ending, unitig index) is sorted by ending, and indexes of unitigs that share the same ending are joined in a tuple. Each such tuple is assigned to a bucket corresponding to one of its unsealed endpoint indices chosen at random (solid arrows in the figure). For the other endpoint index of the tuple (dashed arrow), we put a *placeholder* in its corresponding bucket (in gray). Then, inside each bucket, pairs sharing the same unitig index are joined to form larger tuples. If an ending cannot be joined and does not have a corresponding placeholder, then is marked as sealed and is not selected anymore for bucket assignment. For example, in the first step, in 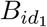 the pair (*id*_1_, *id*_2_) is sealed at *id*_1_, because there is no placeholder for *id*_1_; however, in 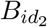 the pair (*id*_2_, *id*_3_) is not sealed at *id*_2_ because *id*_2_ has a placeholder. The steps are repeated until no more tuples can be joined. For the non-canonical case, we can merge the pairs if one extremity is at the end and the other one at the beginning of the pairs. For canonical *k*-mers, we also have to keep track of the direction of the *k*-mer before joining them, see Sec. Construction correctness for more details.

We allocate a fixed number of buckets. Initially, for each list in *P*, we mark both its ends as *unsealed*. We repeat the following procedure until *P* is empty:

1. For each list in *P*, we choose at random one of its unsealed ends. Without loss of generality, let this end be *l*. We put the list in the bucket corresponding to *l*, while in the bucket corresponding to the other ending *r* we put a placeholder.
2. Inside each bucket, we sort the lists by the ending that caused the list to be placed in the bucket. Then, we merge all the endings that are equal to produce longer lists. If an ending in a bucket is not merged, and it has no corresponding placeholder (of another list) in the bucket, then it is marked as *sealed*.
3. Finally, we remove from *P* each list having both ends sealed.

In the above process, given two lists in *P* that must be merged, there is at least a probability of at least 1*/*4 that they are assigned to the same bucket to be merged (in the worst case, both ends are unsealed). Thus, in expectation, this desired outcome happens only after 4 tries.

Both *L* and *P* are also stored in buckets, to allow a better concurrency while processing them.

### Construction correctness

We start by proving the correctness of the algorithm (without reverse complements).

#### Theorem 1.

*Given a multiset R of strings, the strings U obtained at the end of our algorithm are the maximal unitigs of R*.

*Proof*. We prove that *U* satisfied the conditions of Definition 1 (for non-canonical unitigs).

For condition 0, since we start from *k*-mers, all string in *U* have length at least *k*. To see that *U* is also a *set* (i.e., contains no duplicates), in our algorithm every *k*-mer is considered only once, and is assigned to a unique unitig.

For condition 1, observe that the algorithm does not introduce any *k*-mer that is not also in *R*, and does not exclude any *k*-mer from *R*, thus ∀*q* ∈ ∑^*k*^ occ(*q, R*) ≥ 1 iff occ(*q, U*) ≥ 1 holds. By Lemma 1(b), guarantees that occ(*q, R*) the number of occurrence of *k*-mer *q* is the same as the number of occurrences in its group, and thus the operation from Sec. Construction of intermediate unitigs performed inside its group respect its abundance in *R*. Thus, ∀*q* ∈ ∑^*k*^ occ(*q, R*) ≥ *a* iff occ(*q, U*) ≥ 1.

Next, we prove conditions 3 and 2. Given an intermediary unitig *z*, we check for the eight *k*-mers that contain suf_*k*−1_(*z*); we extend *z* iff only two *k*-mers appear (one out-going from, and one in-coming to suf_*k*−1_(*z*)), thus condition 3 is satisfied.

After merging *z* with this out-going *k*-mer, there are no other *k*-mers (and thus no other unitigs in *U*, since each *k*-mer appears exactly once in *U*, because in Sec. Construction of intermediate unitigs we mark the used *k*-mers) that contain suf_*k*−1_(*z*) so condition 2 holds for single groups. This condition is still not satisfied globally, due to the repetition of all the *k*-mers containing a linking character.

We now prove condition 2 after the unitig merging step. Note that the unitigs fed to the unitig merging step are the ones that start or end with a linking character, so they always overlap on *k* characters. After merging all the repeated *k*-mers, we satisfy 2 globally. □

All the steps described so far can be easily adapted to work with canonical *k*-mers, to obtain canonical maximal unitigs (Definition 1). This can be done by changing two steps. First, the hash functions are replaced by their ‘canonical’ version, such that the hash of a *k*-mer is always equal to the one of its reverse complement. Second, in the unitig merging steps, relative orientations of the unitigs are tracked, to allow joining unitigs that can be present in opposite orientations in the input dataset, by reverse-complementing one of them.

### Coloring

Computing the colors for each *k*-mer of a de Bruijn graph has two main challenges: (i) tracking all colors that belong to each *k*-mer, and (ii) storing the colors in a storage- and time-efficient manner. To solve these two challenges, we propose a method partially inspired by the way BiFrost handles the colors, but with numerous improvements that allow for a smaller representation and a faster computation. The main idea is to merge color information for *k*-mers that share the same set of colors, as in [27, 1, 23], while avoiding costly comparisons of the entire sets for each *k*-mer. More precisely, for each *k*-mer, a normalized list *C* of colors is obtained, by tracking the source of each *k*-mer, saving all the colors in a possibly redundant way (for example if the *k*-mer appears multiple times in a reference sequence), and then sorting and deduplicating them. From *C*, a 128-bit strong hash *h* is generated, and is checked against a global hashmap that maps *h* to a color subset index. If a match is not found, then the list *L* is written to the colormap, and a new incremental subset index for *L* is generated. Otherwise, it means that the color set already appeared in a previously processed *k*-mer, so the subset index of that color set is returned. Finally, the *k*-mer in the graph is labeled with its corresponding subset index, which, as discussed above, uniquely identifies a color subset. Overall, this allows a better compression, since each subset is encoded only once and not for every *k*-mer that belongs to it.

To optimize the disk space of the color map, this is encoded using a run-length compression scheme on the differences of the sorted colors of the subset, then it is divided into chunks for faster access and compressed again with a run of the lz4 algorithm. Furthermore, when writing the unitigs to disk, we mark the colors of each unitig in the header of the unitig sequence in the fasta file, by also run-length encoding the color set indices of all the *k*-mers of the unitigs. This strategy works well because most unitigs are “variation-free”, and thus tend to have only a small number of possible color subsets associated to its *k*-mers.

### Sequence querying

GGCAT performs queries by dividing unitigs of the input graph and the queries in buckets, using an approach similar to the reads splitting step of the build algorithm. Then, independently for each bucket, a *k*-mer counting is done to find the number of *k*-mers that match for each query. Finally all the counters from different buckets are summed up to find for each query the number of *k*-mers that are present in the input graph. This allows also the partial matching of queries, since the output is the exact number of *k*-mer matches for each input sequence, and a percentage of required matching *k*-mers can be put as threshold to report a query as present. Similarly to BiFrost, for the uncolored case, we return in output a .csv file with a line for each input query, containing the number and percentage of matched *k*-mers. For the colored case, we opted instead for a JSON Lines (.jsonl) file with a line for each query, containing the number (if positive) of *k*-mer matches for each color *c* of the graph.

### Data access

GGCAT is implemented in Rust and is available at GitHub (https://github.com/algbio/ggcat). The Human Illumina read dataset is the Illumina WGS 2X250bp dataset from the GIAB project, accession number HG004, https://github.com/genome-in-a-bottle/giab_data_indexes/blob/master/AshkenazimTrio/sequence.index.AJtrio_Illumina_2x250bps_06012016.HG004. The Human gut microbiome read dataset is from project PRJEB33098 [24]. The 309K Salmonella genome sequences were downloaded by us in February 2022 from the EnteroBase database [38] and gzipped. The 100 Human genomes are from the set of 2505 Human genomes generated by [34] using GRCh37 and the variant files from the 1000 Genomes Project [36]. For convenience we uploaded the Human genomes used for the benchmark to Zenodo (10.5281/zenodo.7506049, 10.5281/zenodo.7506425). The Bacterial genome sequences are from the dataset used in [3].

For all tests we used GGCAT commit f56da4d35f99f3537ec0a33f44d575898a8c91ea (https://github.com/algbio/ggcat/tree/f56da4d35f99f3537ec0a33f44d575898a8c91ea). The tools and scripts used to perform the benchmarks are available at https://github.com/Guilucand/ggcat-test-benchmarks. The scripts to download the tests data are available in the datasets-download/ folder of the above repository.

## Competing interests

The authors declare no competing interests.

## Acknowledgements

We are grateful to Massimo Cairo and Romeo Rizzi for initial discussions on this problem. We are grateful to Paul Medvedev for spotting some errors in the initial manuscript, pointing us to relevant literature, and suggesting that the equivalence between edge-centric and node-centric unitigs exists, which we proved in the Supplemental Material. We thank Rob Patro and Jarno Alanko for pointing us to relevant literature. We thank Sebastian Schmidt for help with the integration of eulertigs [35] and matchtigs [34] into GGCAT.

This work was partially funded by the European Research Council (ERC) under the European Union’s Horizon 2020 research and innovation programme (grant agreement No. 851093, SAFEBIO), and partially by the Academy of Finland (grants No. 322595, 352821, 346968).

## Supplemental Material

### 1 Equivalence of node-centric and edge-centric unitigs

In this section we prove the equivalence between the (maximal) unitigs of the node-centric de Bruijn graph and the (maximal) unitigs of the edge-centric de Bruijn graph, built on the same set of strings, *R*, (i.e., their spellings are exactly the same strings). Note that we give this proof only for directed graphs.

For ease of notation, in this section we will denote the edge-centric graph of *R* as 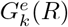. The *node-centric* graph for *R*, which we denote as 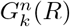, is formally defined by adding a node for every *k*-mer of *R*, and an edge between two nodes *x* and *y* if suf_*k*−1_(*x*) = pre_*k*−1_(*y*). In a node-centric graph, a path *P* (containing at least one node) is said to be a *node-centric unitig* if all nodes of *P*, except the last node, have out-degree equal to one, and all nodes of *P*, except the first one, have in-degree equal to one [1]. A node-centric unitig is said to be *maximal* if it cannot be extended by a node on either side [1]. See Figure 1 for an example. We will use the term *unitig* to refer to a unitig in the edge-centric graph (as defined in the Preliminaries subsection of the main paper), and *node-centric unitig* to refer to the unitigs just defined in the node-centric graph.

**Figure 1:**
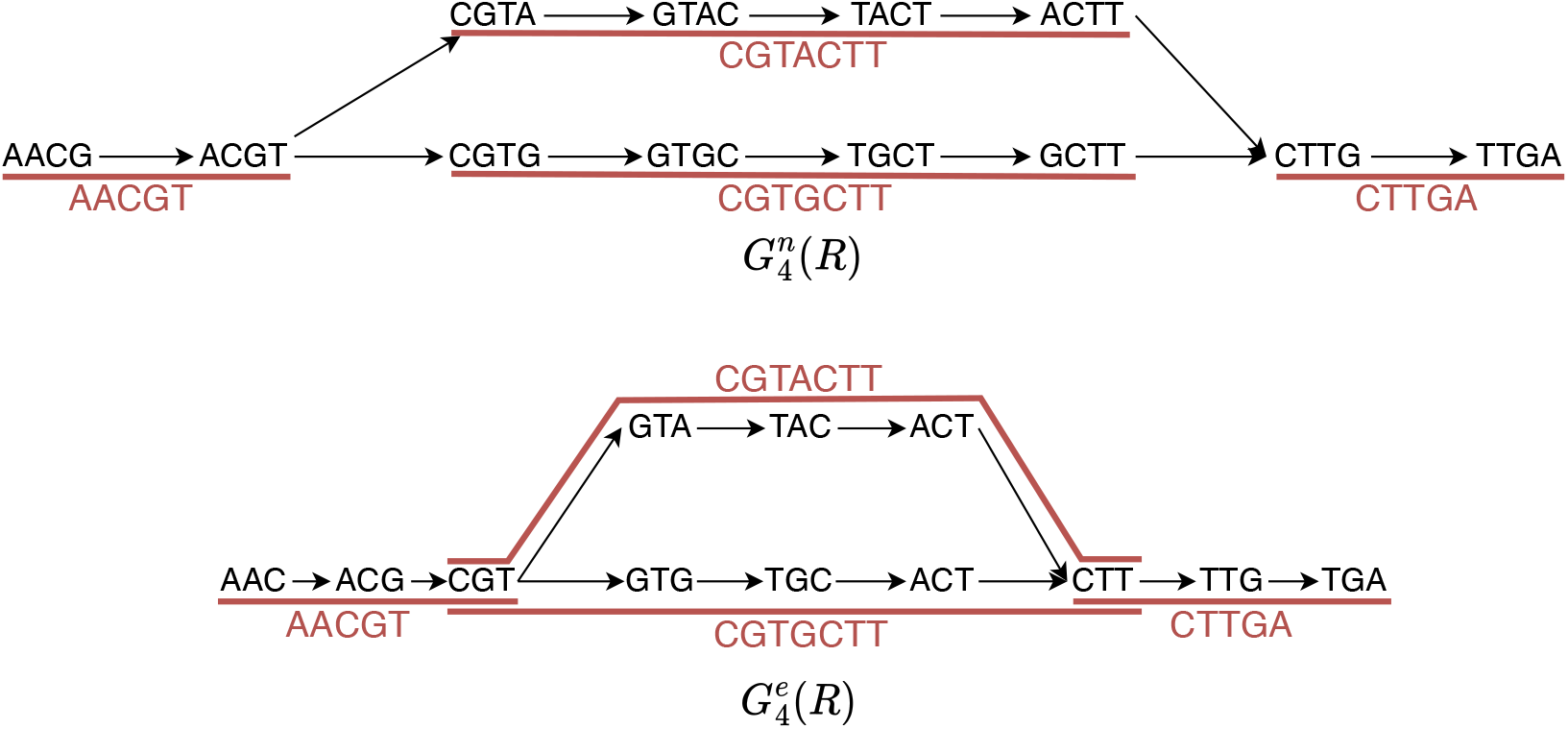
Top: Illustration of the node-centric de Bruijn graph 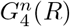, where we assume *R* is the set consisting of all 4-mers that label the nodes of the shown graph. Bottom: the edge-centric de Bruijn graph 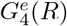, built on the same set *R*. In both graphs we draw in red their corresponding maximal unitigs (i.e., node-centric and edge-centric, respectively). In the node-centric graph, the central unitigs do not include the node ACGT, since its out-degree is different from one, and do not include the node CTTG since its in-degree is different from one. Every unitig is labeled with its spelling, also in red.

The *spelling* of a path *P* = (*x*_1_, …, *x*_*t*_) in 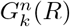 is analogously defined as the string *x*_1_ ⊙^*k*−1^ *· · ·⊙*^*k*−1^ *x*_*t*_. As in the edge-centric case, by a node-centric unitig we will refer to either a path *P* in the node-centric graph, or to the spelling of *P*.

#### Theorem 1.

*Let R be a set of strings, and let 𝒳 be the set of all node-centric unitigs of* 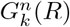 *and let 𝒴 be the set of all unitigs of* 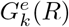. *Then 𝒳* = 𝒴.

*Proof*. We prove the claim by proving both implications. See Figure 2 for an illustration of the two analogous cases.

(𝒳 ⊆ *𝒴*): Let *X* ∈ *𝒳*, we show that *X* ∈ *𝒴* holds. Suppose *X* = (*x*_1_, …, *x*_*p*_), and let *y*_*i*_ := pre^*k*−1^(*x*_*i*_), for *i* ∈ *{*1, …, *p}*, and *y*_*p*+1_ := suf_*k*−1_(*x*_*p*_). Since *x*_1_, …, *x*_*p*_ are *k*-mers of *R*, then the edges (*y*_*i*_, *y*_*i*+1_), *i* ∈ *{*1, …, *p}*, exist in 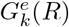. We claim that *Y* := (*y*_1_, …, *y*_*p*+1_) is a unitig in 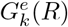 (which clearly has the same spelling as *X*). First, note that *Y* contains at least one edge, since *p* ≥ 1. Suppose for a contradiction this is not the case, namely that there is some *y*_*i*_, *i* ∈ *{*2, …, *p}*, having w.l.o.g. out-degree different than one. Since the edge (*y*_*i*_, *y*_*i*+1_) exists in 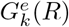, this means that the out-degree of *y*_*i*_ is non-zero, and thus at least two. Let (*y*_*i*_, *y*^*′*^) be another edge out-going from *y*_*i*_ (*y′* ≠ *y*_*i*+1_). Therefore, *x′* := *y*_*i*_ *⊙* ^*k*−2^ *y′* is a node in 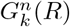. Moreover, since *i* ≥ 2, there is a node *y*_*i*−1_ preceding *y*_*i*_ in the unitig *Y*, and thus a node *x*_*i*−1_ preceding *x*_*i*_ in the unitig *X*. Consider now the nodes *x*_*i*−1_, *x*_*i*_, *x′* in 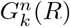. By definition, we have that suf_*k*−1_(*x*_*i*−1_) = pre^*k*−1^(*x*_*i*_), and suf_*k*−1_(*x*_*i*−1_) = *y*_*i*_ = pre^*k*−1^(*x*^*′*^). Thus, the out-degree of *x*_*i*−1_ in 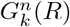 is at least two, which contradicts the definition of node-centric unitig, since obviously *x*_*i*−1_ is not the last node of the unitig *X*.

(𝒴 ⊆ *𝒳*): Let *Y* ∈ *𝒴*, we show that *Y* ∈ *𝒳* holds. Suppose *Y* = (*y*_1_, …, *y*_*p*_), *p* ≥ 2, and let *x*_*i*_ := *y*_*i*_ *⊙*^*k*−2^ *y*_*i*+1_, *i* ∈ *{*1, …, *p* − 1*}*. Since *x*_1_, …, *x*_*p*_ are (*k* − 1)-mers of *R*, then they are all nodes in 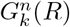, and *X* := (*x*_1_, …, *x*_*p*−1_) (which has the same spelling as *Y*) is a path in 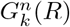 (note that *p* − 1 ≥ 1). We claim that *X* is a node-centric unitig in 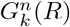. Suppose for a contradiction that this is not the case. Assume that there is *x*_*i*_, *i* ∈ *{*1, …, *p* − 2*}*, having w.l.o.g. out-degree different from one, and thus at least two. Let *x′ ≠ x*_*i*+1_ be another out-neighbor of *x*_*i*_, and let *y′* := suf_*k*−1_(*x′*). Consider now the nodes *y*_*i*+1_, *y*_*i*+2_, *y′* (recall *i* ≤ *p* − 2) in 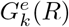. By definition, (*y*_*i*+1_, *y*_*i*+2_) is an edge in 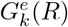; moreover (*y*_*i*+1_, *y*^*′*^) is also an edge, because *x′* = *y*_*i*+1_ ^*k*−2^ *y′* is a node in 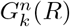. Thus, the out-degree of *y*_*i*+1_ is at least two. Since *i* ≤ *p* − 2, this means that *y*_*i*+1_ is a node of the unitig *Y*, different from its last node, having out-degree at least two, which is a contradiction.

Next, we prove the same equivalence, but for *maximal* unitigs.

**Figure 2:**
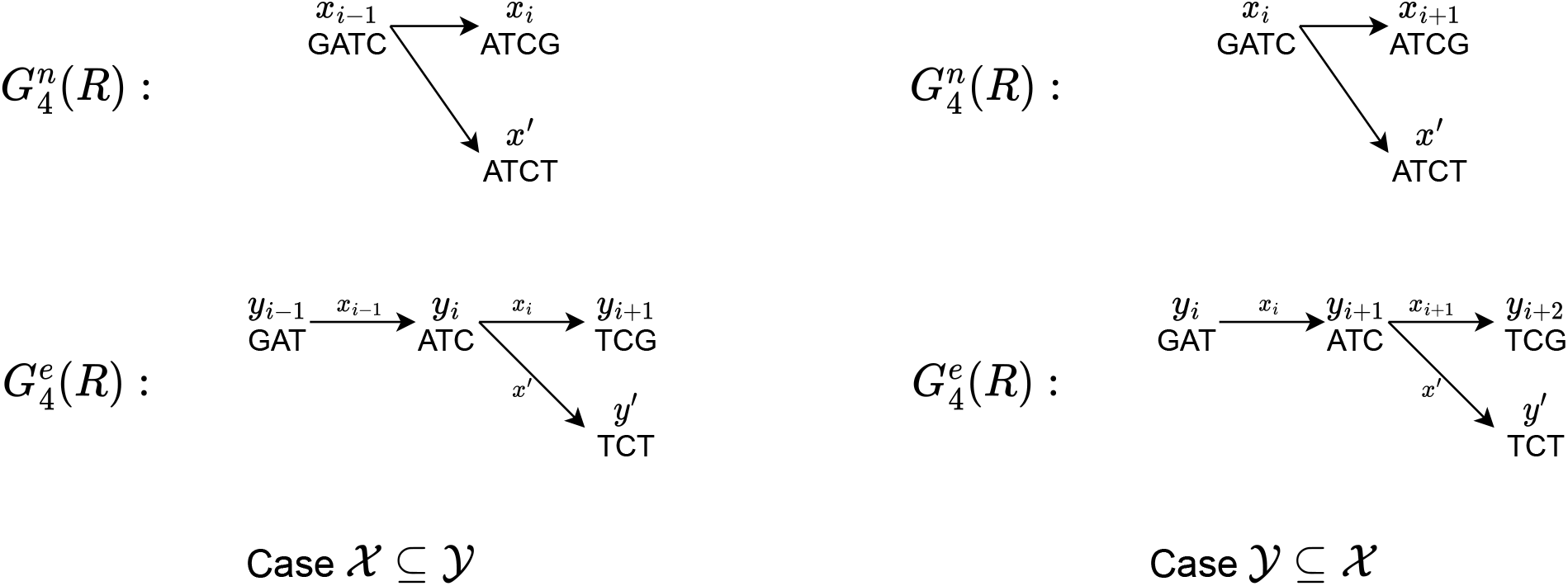
Illustration of the two analogous cases in the proof of Theorem 1. For concreteness, we also draw some example strings on the nodes of graphs. In the edge-centric graphs, we label the edges with their corresponding *k*-mer.

#### Corollary 1.

*Let R be a set of strings, and let 𝒳 be the set of all maximal node-centric unitigs of* 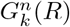 *and let 𝒴 be the set of all maximal unitigs of* 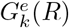. *Then 𝒳* = 𝒴.

*Proof*. We prove one inclusion only, since the other one follows completely symmetrically. Let *M* be a maximal node-centric unitig in 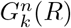. We want to prove that *M* is also a maximal unitig in 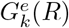. Suppose for a contradiction that *M* is not maximal in 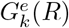. Then there exists a unitig *M′* of 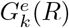, *M* ∈ *M′* with |*M′* | *>* |*M* |. By Theorem 1, we have that *M* is also a node-centric unitig in 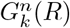. Since *M* ∈ *M*^*′*^, this contradicts the maximality of *M* in 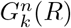. Thus, *M* is a maximal unitig of 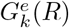. □

## 2 Supplementary results

**Figure 3:**
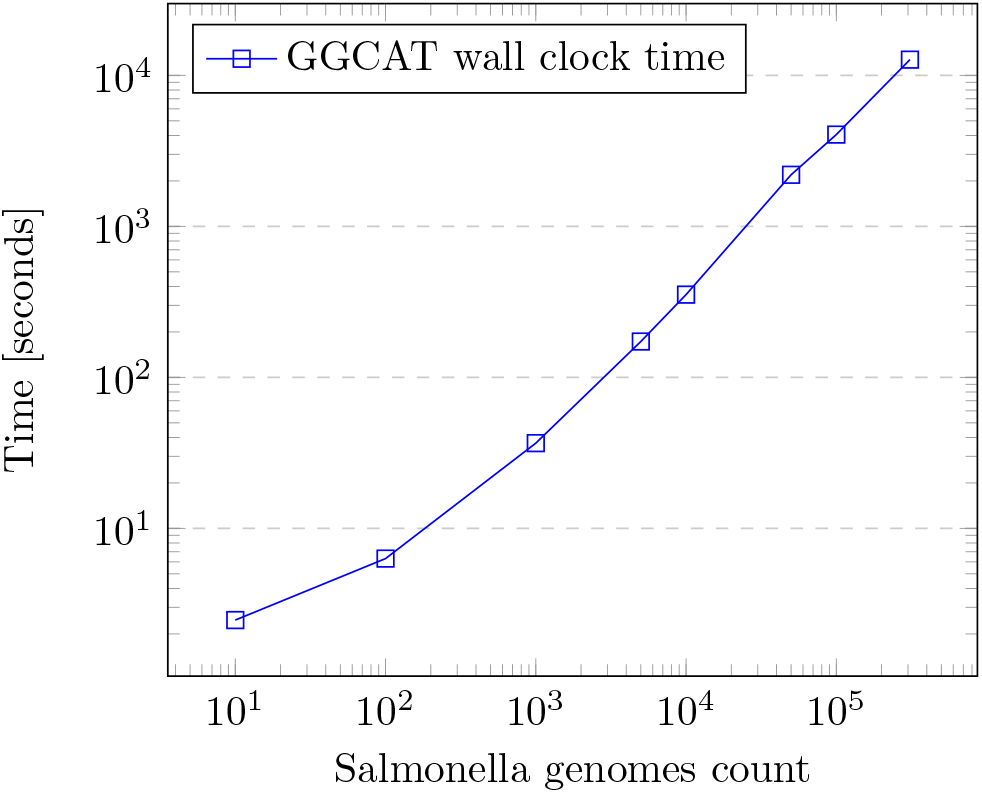
GGCAT running time with an increasing amount of Salmonella genomes, *k* = 63, using 12 threads, on the small server.

## 3 Commands used

Here we list all the command templates used to benchmark the tools.

### 3.1 Uncolored building

~~~
# Cuttlefish 2 for sequencing reads
./cuttlefish build --read -l <INPUT_FILES_LIST> -k <KVALUE> -c <MULTIPLICITY> -t <THREADS> -o <OUTPUT_FILE> -w <TEMP_DIR>
# Cuttlefish 2 for reference genomes
./cuttlefish build --ref -l <INPUT_FILES_LIST> -k <KVALUE> -c <MULTIPLICITY> -t <THREADS> -o <OUTPUT_FILE> -w <TEMP_DIR>
# GGCAT for both reads and reference genomes
./ggcat build -k <KVALUE> -j <THREADS> -s <MULTIPLICITY> -l <INPUT_FILES_LIST> -t <TEMP_DIR> -o <OUTPUT_FILE>
~~~

### 3.2 Colored building

~~~
# Bifrost
./Bifrost build -k <KVALUE> -t <THREADS> <INPUT_FILES_LIST> -o <OUTPUT_FILE> --verbose -c
# GGCAT
./ggcat build -k <KVALUE> -j <THREADS> -s <MULTIPLICITY> -l <INPUT_FILES_LIST> -t <TEMP_DIR> -o <OUTPUT_FILE> --colors
~~~

### 3.3 Colored querying

~~~
# Bifrost
./Bifrost query -k <KVALUE> -t <THREADS> -g <INPUT_GRAPH> -q <INPUT_QUERY> -o <OUTPUT_FILE> --verbose
# GGCAT
./ggcat query -k <KVALUE> -j <THREADS> <INPUT_GRAPH> <INPUT_QUERY> -t <TEMP_DIR> -o <OUTPUT_FILE> --colors
~~~

## Footnotes

1 In this paper we use the edge-centric definition of a de Bruijn graph, and hence the edge-centric version of the notion of unitig, see Sec. Preliminaries.

2 Since the first and the last *k*-mer of every maximal unitig is a node of the compacted de Bruijn graph, it often suffices to compute only the strings labeling the maximal unitigs of a de Bruijn graph, not the actual graph structure. Moreover, reverse complements need special handling, which we describe in Sec. Preliminaries.

3 Our definition of unitigs is for edge-centric graphs. For *node-centric de Bruijn graphs*, unitigs need to be defined by imposing an additional condition, see e.g. [9, 16]. To the best of our knowledge, we are not aware of a formal proof of equivalence between these types of unitigs in the two types of graphs. As such, we give this proof in the Supplemental Material.

4 In this paper, paths do not repeat nodes, except for possibly, the first and the last, in which case we say that the path is a *cycle*. If two cycles *C*_1_ and *C*_2_ are such that *C*_2_ can be obtained from *C*_1_ by a cyclic shift, then we say that *C*_1_ and *C*_2_ are *equivalent*.

## Notes

https://github.com/algbio/ggcat

### Competing Interest Statement

The authors have declared no competing interest.

### Summary of Updates

Updated citations for colored construction methods Added new bacterial genomes experiments Added data access section Updated supplemental files with proof of edge and node based de Bruijn graphs equivalence

